# Transversal otolithic membrane deflections evoked by the linear accelerations

**DOI:** 10.1101/844977

**Authors:** V. Goussev

## Abstract

Considered is the model of the transversal utricle membrane deflections evoked by the linear accelerations. The real 3D utricle membrane structure was simplified by considering its middle section and evaluating its elastic properties in 2D space. The steady state transversal deflections along the membrane are analytically evaluated and numerically simulated using the 2D elasticity theory. The transversal deflections are found to be more expressive and stronger as compared to the conventional longitudinal deformations. The revealed properties could be used for explanation of the transduction processes in the otholith organ. Based on the implemented modeling approach the new otolithic membrane mechanical properties are discussed and new explanations for the available experimental data are given.

## Introduction

The otolith organ in the vestibular system plays an essential role being the sensor of linear accelerations. Many researchers (Igarashi, 1966; Twizell & Curran, 1977; Kondrachuk, 2001; Li, Xue, & Peterson, 2008) in the past and nowadays studied the information conversion properties of the otolith organ both in the physiology and biophysics. The otolith organ consists of two independent sensors: utricle and sacculus, containing both otolithic membranes with different layers and the hair cells, which are sensitive to the membrane deflections and, hence, to the linear accelerations. The membranes look like the enclosed, almost flat capsules, which are rigidly attached to the bones at all their boundaries, except the utricle, which is attached to the bones only at its longest end (Uzun-Coruhlu, Curthoys & Jones, 2007). However, its attachment to the supporting flexible membrane in other boundary regions can be also considered as rigid, as the supporting membrane is more solid than the utricle capsule since it contains the hair cell bodies and afferent and efferent fibers. The surface of otolithic membranes is not perfectly flat and has a complicated shape in 3D space. However, some researchers considered them as fully flat (Hudetz, 1973; McGrath, 2003) or slightly spherical (Jaeger, 2003). The otolithic membranes of utricle and sacculus are approximately orthogonal in space, what helps them decomposing the arbitrary linear accelerations into projections of the body reference systems. Both utricle and sacculus sensors are similar in their transduction properties, so we shell concentrate further on the transduction properties of the utricle.

### Transition from 3D to 2D space

From the point of view of the elasticity theory the real utricle membrane can be considered as the plate with the varying curvature in 3D space. Because of the apparent complexity, the most reliable and convinced way to study its transduction properties is the numerical 3D simulation, realized usually with the finite element method (Sato, Sando & Takahashi, 1992; Kondrachuk, 2001; Jaeger, 2003; Silber et al.,2004). However, to be able to get some preliminary analytic results, we would like to simplify the problem considering only the fraction of the utricle membrane, extracting the strip in the middle of the membrane along its longest axis. The extracted strip of the arbitrary width can be considered as a thin beam simply supported at its end and slightly spherically curved. The extraction of the thin beam from the homogeny region of the utricle membrane doesn’t change its elastic property as compared with the whole membrane, but helps implementing 2D elastic theory (Timoshenko & Gere, 1961) to its deflection analysis.

### Deflection of the utricle membrane

The beam cut out from the middle region of the utricle preserves its complex structure represented on the Figure 1. The structure consists of three layers: otoconia, mesh, and gel like fluid. The geometrical values for all membrane layers vary from sources. In most cases we shall follow the geometrical data presented in Jaeger’s work (Jaeger, 2003). Our specific attention is paid for the clearly demonstrated cavities in the mesh layer (Kachar, Parakkal & Flex, 1990), which are filled out with a gel like fluid and cover the hair cell bundles, giving impression that the mesh layer is perforated.

**Figure 1.**
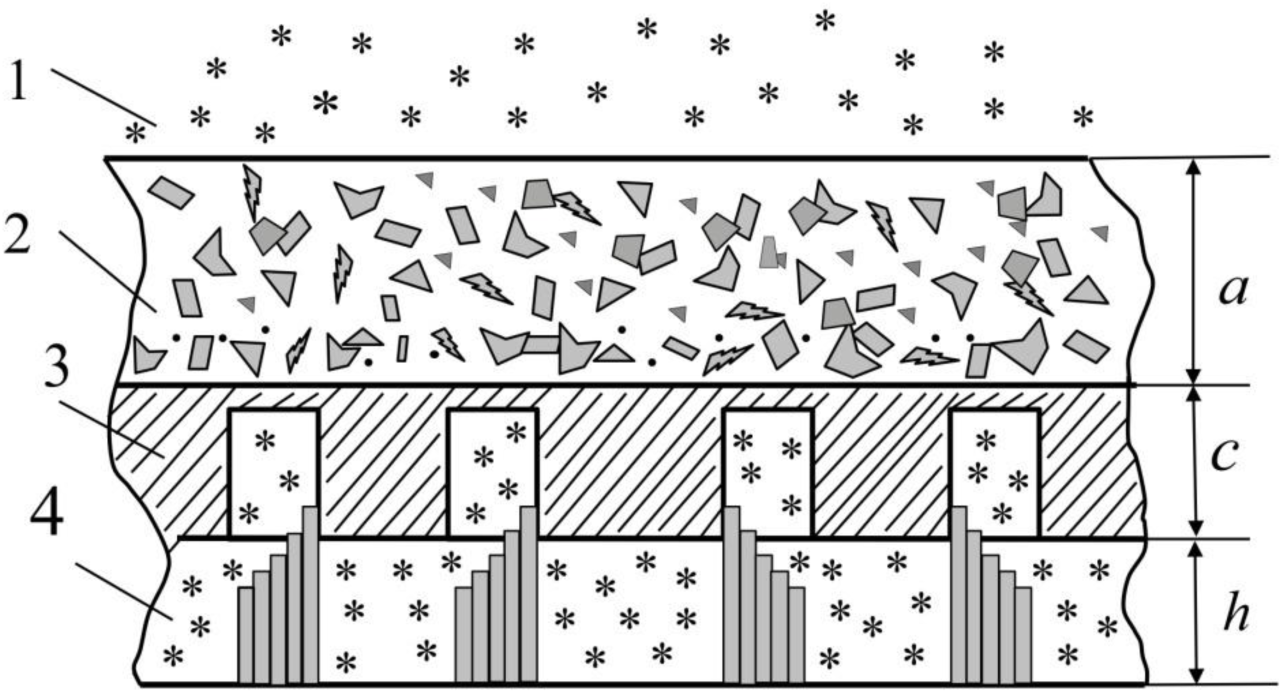
Schematic fragment of the utricle membrane and hair cells. 1-endolymph, 2-otoconia layer, 3-mesh layer, 4 – gel layer. The mean geometrical values: a=15 mkm, c=10 mkm, h=10 mkm.

The basic idea underlying our consideration is that the longitudinal load may cause the utricle membrane deflections, bending it in the buckling way. We consider the replacement of the membrane fragment (Figure 1) by the conventional simply supported beam of the constant rectangular cross section, which loading scheme is represented in Figure 2. The beam is loaded by the homogeneous longitudinal force, which is evoked by the linear acceleration. It is supposed that the beam is associated mainly with the otoconia layer, as it has the most rigid structure as compared with other layers. The height of the beam is 15 mkm and its length is assumed to be about 1000 mkm. The width of the beam cross section is not defined in the 2D elasticity problem and can be assumed to be 1. The beam is also assumed to be slightly spherically curved with the curvature radius of 10 mm. The most uncertain parameter is the elasticity modulus E (Young modulus), which is varying essentially from sources from 2000 Pa (Jaeger,2003) to 6.6 MPa (<u>Davis, Xue, Peterson & Grant</u>, 2007). We also evaluate it later based on our physical assumption that the elastic properties of the beam are defined entirely by the available range of the membrane deflections.

**Figure 2.**
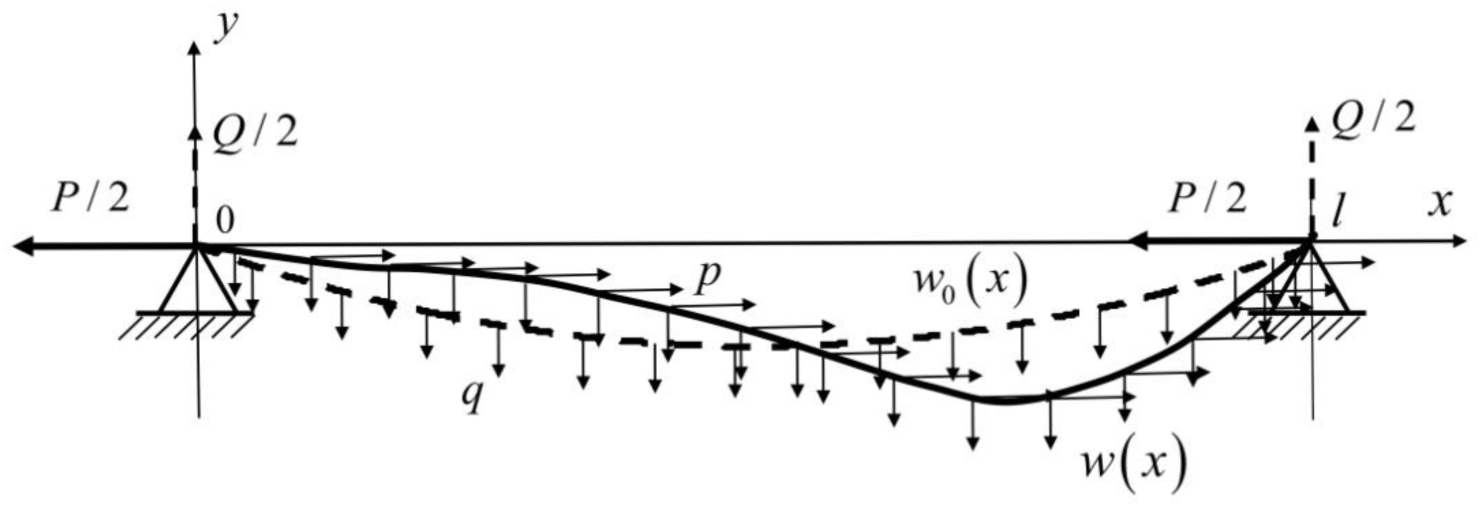
The utricle otoconia layer under the acceleration buckling load.

The beam is assumed to be initially deflected *w*_0_ (*x*) by the fictitious transverse load *q*, allowing to imitate the initial beam shape. Additional longitudinal load *p* deflects the beam to its steady state shape *w*(*x*). The forces *P* / 2 and *Q* / 2 are the support forces required for the static stability. All three membrane layers follow the initial beam shape *w*_0_ (*x*), so the gel layer height *h* is approximately constant along the beam. During the additional longitudinal loading only the mesh layer follows the deflection of the otoconia layer, while the gel like fluid is forced to flow along the layer space changing its local height. Hence, the difference Δ*w* = *w*(*x*)− *w*_0_ (*x*) is the deflection of the beam relative its initial position and in the same time is the change in the gel layer height. As the total volume of the gel like liquid should be constant in the utricle capsule, the liquid should spill over from the region with the high pressure to the lower pressure regions along the membrane, so the following restriction should take place:

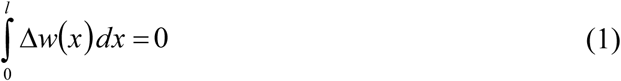

The beam deflection *w*(*x*) can be derived from the balance of works made by internal and external forces (Timoshenko & Gere, 1961):

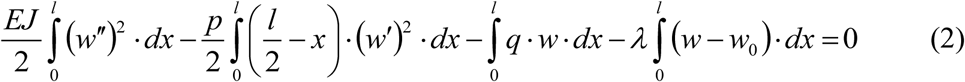

where *E* - the elasticity module, *J* - the moment of inertia of the beam cross section, *λ* - the Lagrange multiplier, *l* - the length of the beam.

The balance of the works (2) is reached when the first variation of Equation (2) relative to the arbitrary variation *δw* of deflection *w*(*x*) is getting zero. The resulting equation for evaluation of deflection *w*(*x*) reads:

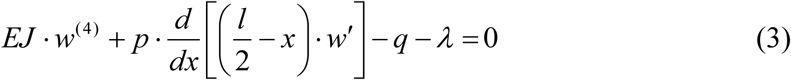

Taking into account that the initial deflection *w*_0_ (*x*) can be considered as the solution of the equation:

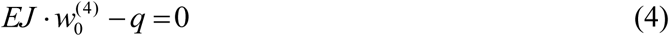

we obtain the part of deflection Δ*w*, which is evoked by the longitudinal load, from the difference between Equations (3) and (4):

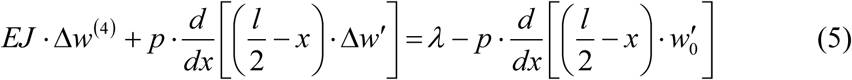

Integrating Equation (5) results:

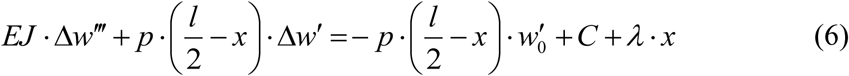

where *C* and *λ* are arbitrary constants.

Denoting *u* = Δ*w′* we get the simplified version of the Equation (6):

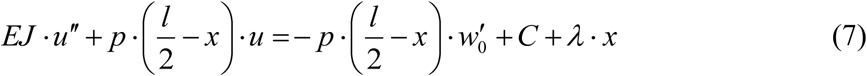

with boundary conditions: 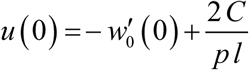, *u′*(0) = 0. Constants *C* and *λ* can be evaluated from the boundary conditions at the end of the beam: Δ*w*(*l*) = 0 and the restriction (1).

### Numerical simulation

Equation (7) is the non-homogeneous linear differential equation with the space varying coefficient. One of the most convenient method to get to the solution of Equation (7) is its replacement by the matrix equation. Denoting 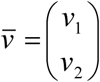, *v*_1_ = *u* and *v*_2_ =*u′* we can rewrite Equation (7) in the matrix form:

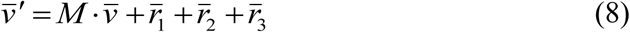

where 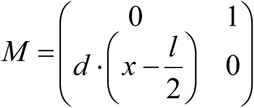, 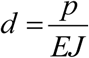, *p* = *γρab*, *γ* is the linear acceleration, *a* is the height of the otoconia layer, *b* is the beam width (assumed to be 1 in the 2D elasticity problems), *ρ* is the otoconia layer density. The external inputs 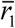, 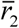, 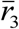 are vectors: 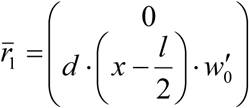, 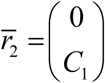, 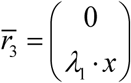, where 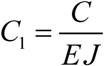, 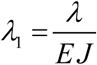. The numerical procedure to get to the solution of the Equation (8) was implemented in Matlab on the grid (2,1001) for the beam length *l* =1*mm* with the spacing Δ*x* = 0.001 *mm*. The Equation (7) was solved separately for every inputs 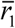, 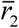, 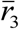. Starting with the initial values *C*_10_ = 1, *λ*_10_ = 1 and the elasticity module *E* = 10^4^ *Pa* the solutions 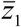, 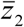, 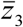 with the initial conditions: 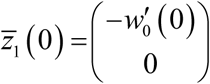, 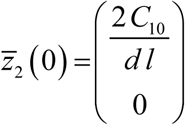, 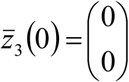 were integrated in *x* ∈[0,*l*] and used then to form the resulting beam deflection Δ*w* as a weighted sum fitting the boundary condition *Δw*(*l*) = 0 and the restriction (1).

## Results

First, we illustrate the influence of the membrane curvature on the otoconia layer deflection changing the radius of curvature *R* in the range (3.6-10) mm via equation: *R*_*i*_ =10 / (1+ (*i* −1)/ 5), *i* =1,…,10. For the linear acceleration *γ* = *g* / 2, *ρ* = 2300 *kg* / *m*^3^ and *E* =10^4^ *Pa* the set of deflections of the otoconia layer is represented on Figure 3.

**Figure 3.**
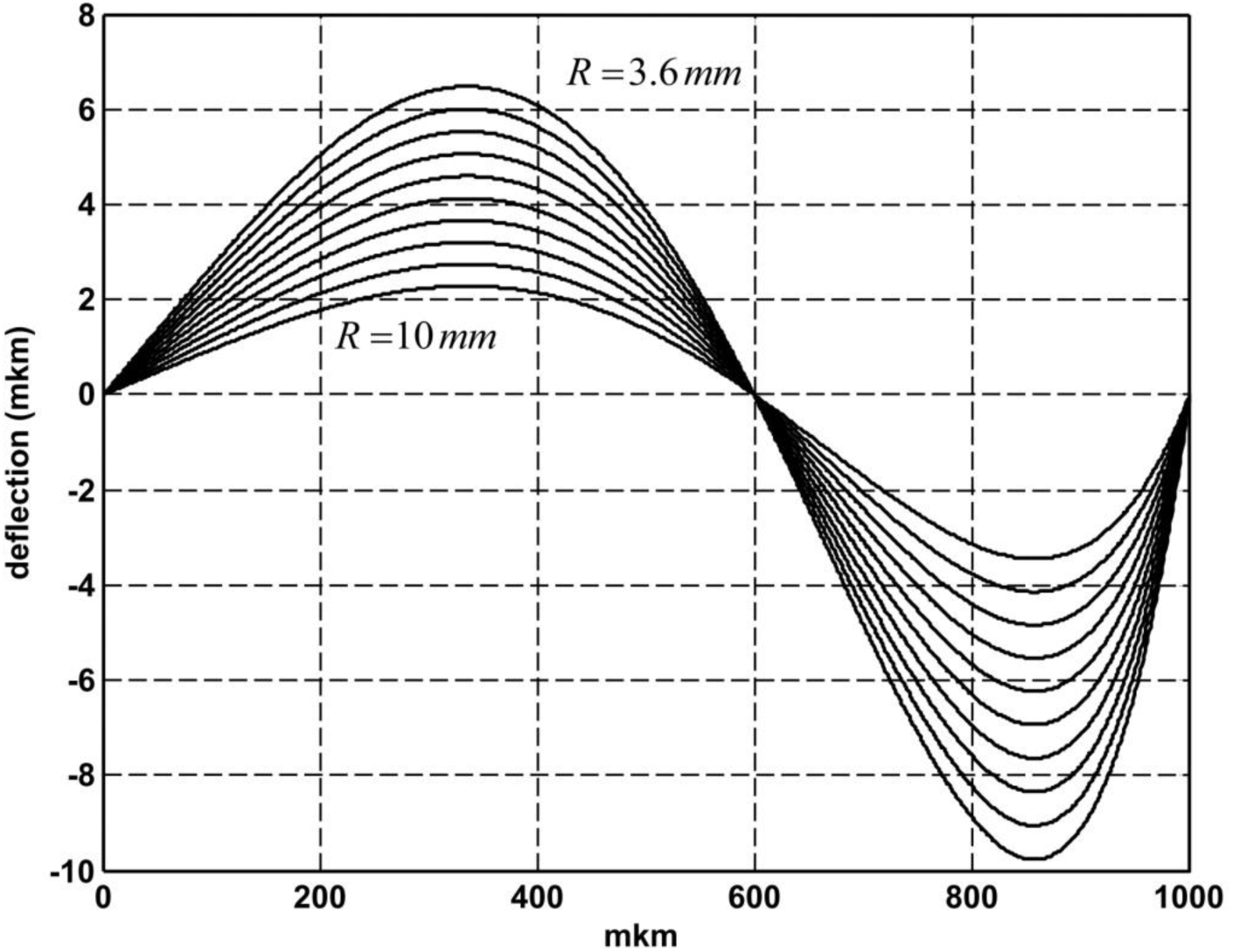
The utricle membrane deflections for the set of its initial curvature radius. The radius is equally distributed between values from 3.6 mm to 10 mm.

It is of the main interest to see the basic transduction property of the utricle membrane deflection as the response to the input linear accelerations. Starting with the threshold value of 0.01g (Kingma, 2005) we limit us with 1g as the highest value typical for the conventional life conditions. For the set of the linear accelerations given by equation: *γ*_*i*_ = *g* / *i*^2^, *i* =1,…,10 and *E* =10^4^ *Pa* the set of the otoconia layer deflections is represented on the Figure 4. It could be seen from the Figure 4 that the whole range of the input acceleration values fits the range of the possible height changes in the gel layer. This evidence was used to evaluate the elasticity modulus as *E* =10^4^ *Pa*. Using lower values leads to the sticking of the mesh layer to the floor region of the gel layer, whereas higher values result in small deflections. The values of deflections as the response to the linear accelerations are essentially non linear. To be able to see all details of deflections we need an additional scaling for the deflections resulting from the low values of accelerations. This was achieved by the scale factor *c*, which dependence on the acceleration values is represented on the Figure 5.

**Figure 4.**
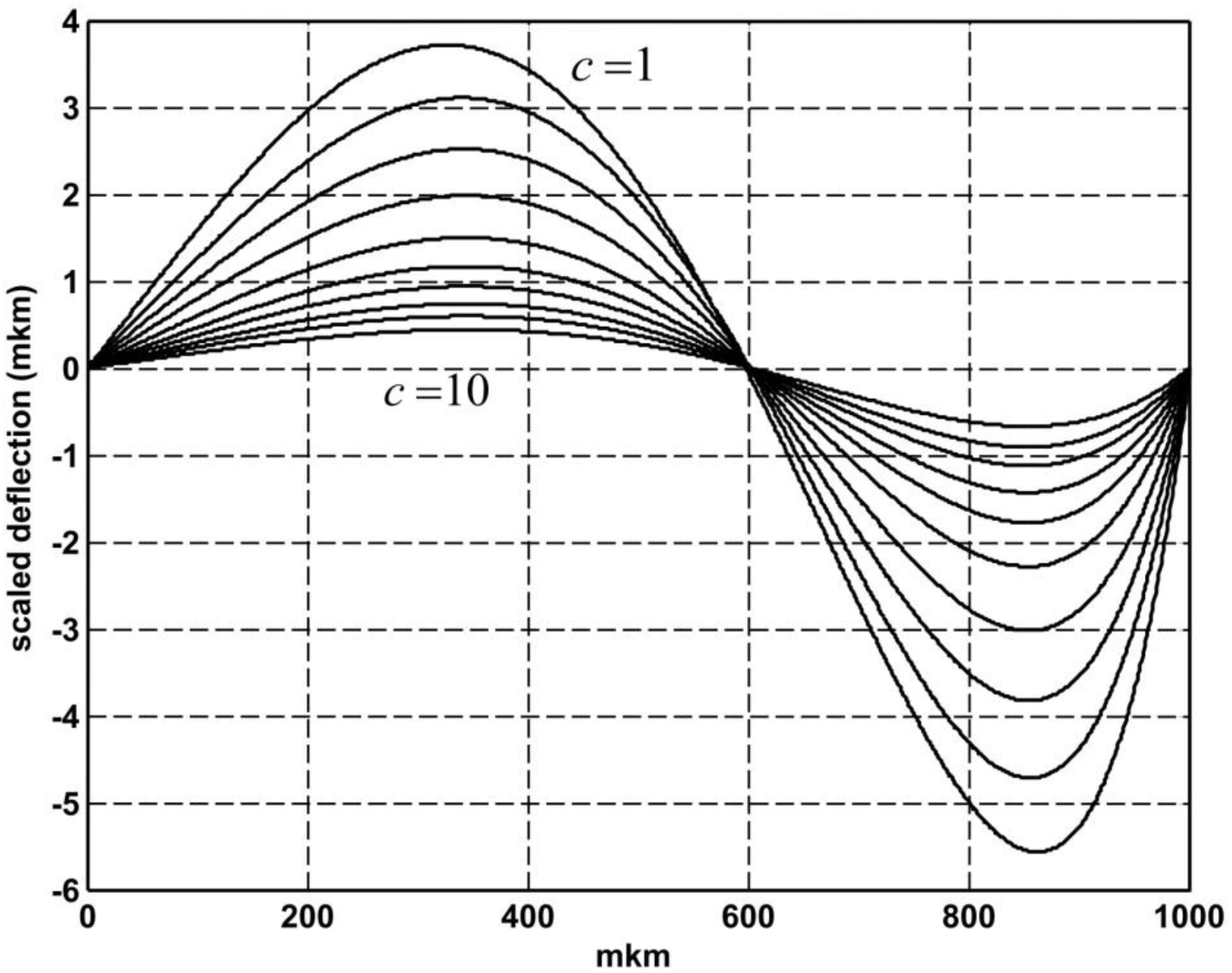
The scaled utricle membrane deflections for the linear acceleration set.

**Figure 5.**
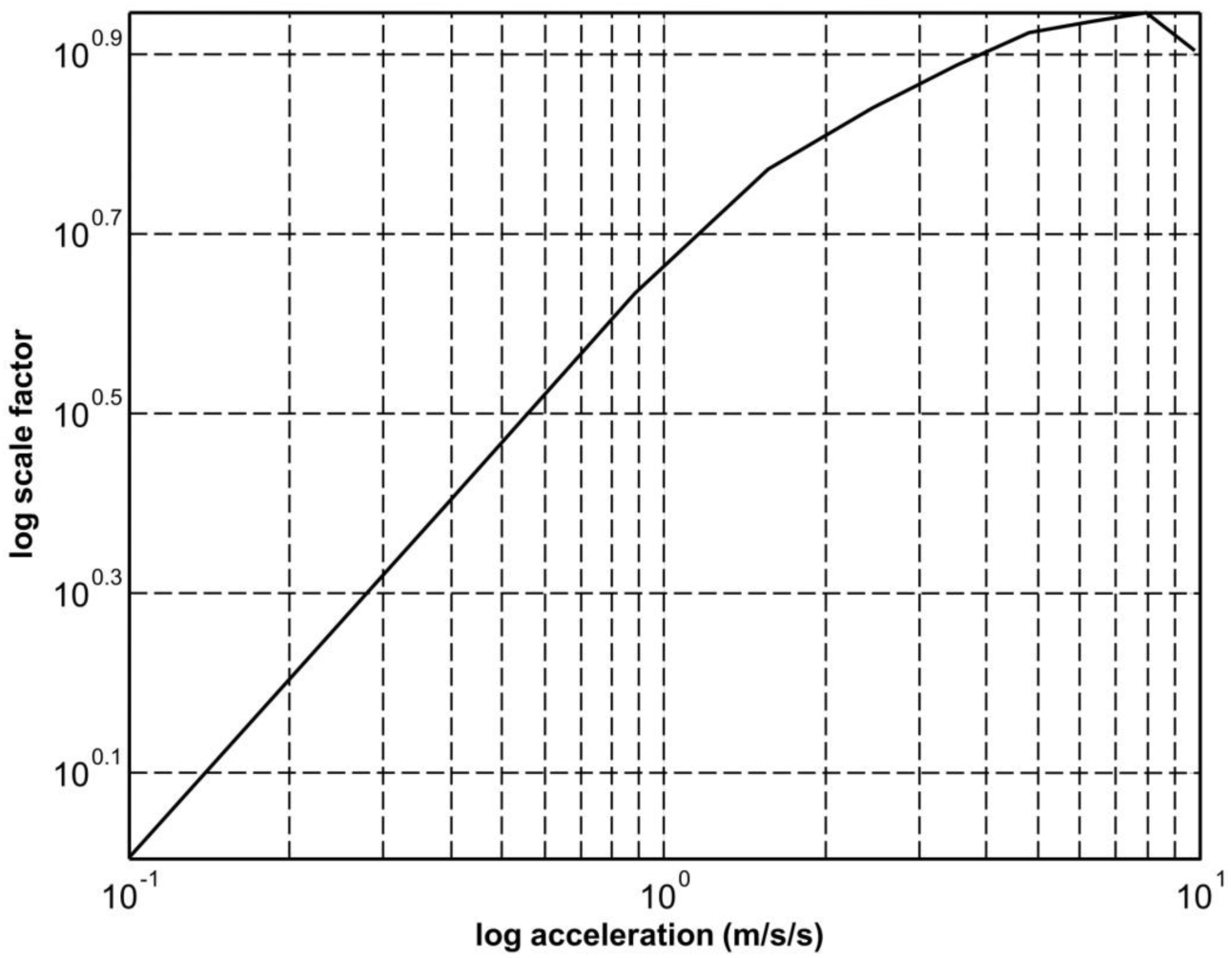
The scaling factors for displaying Figure 4

Instead of the separate scaling for the every otoconia layer deflection given on the *y* axis in Figure 4, we can plot the same set of deflections with the unified scale on the logarithm axis *y*_1_. This scale of the deflection axis is made by the function: *y*_1_ = *sign*(*y*)⋅log(1+8⋅10^6^ ⋅*abs*(*y*)). The new scaled set of the otoconia layer deflections is shown in the Figure 6.

**Figure 6.**
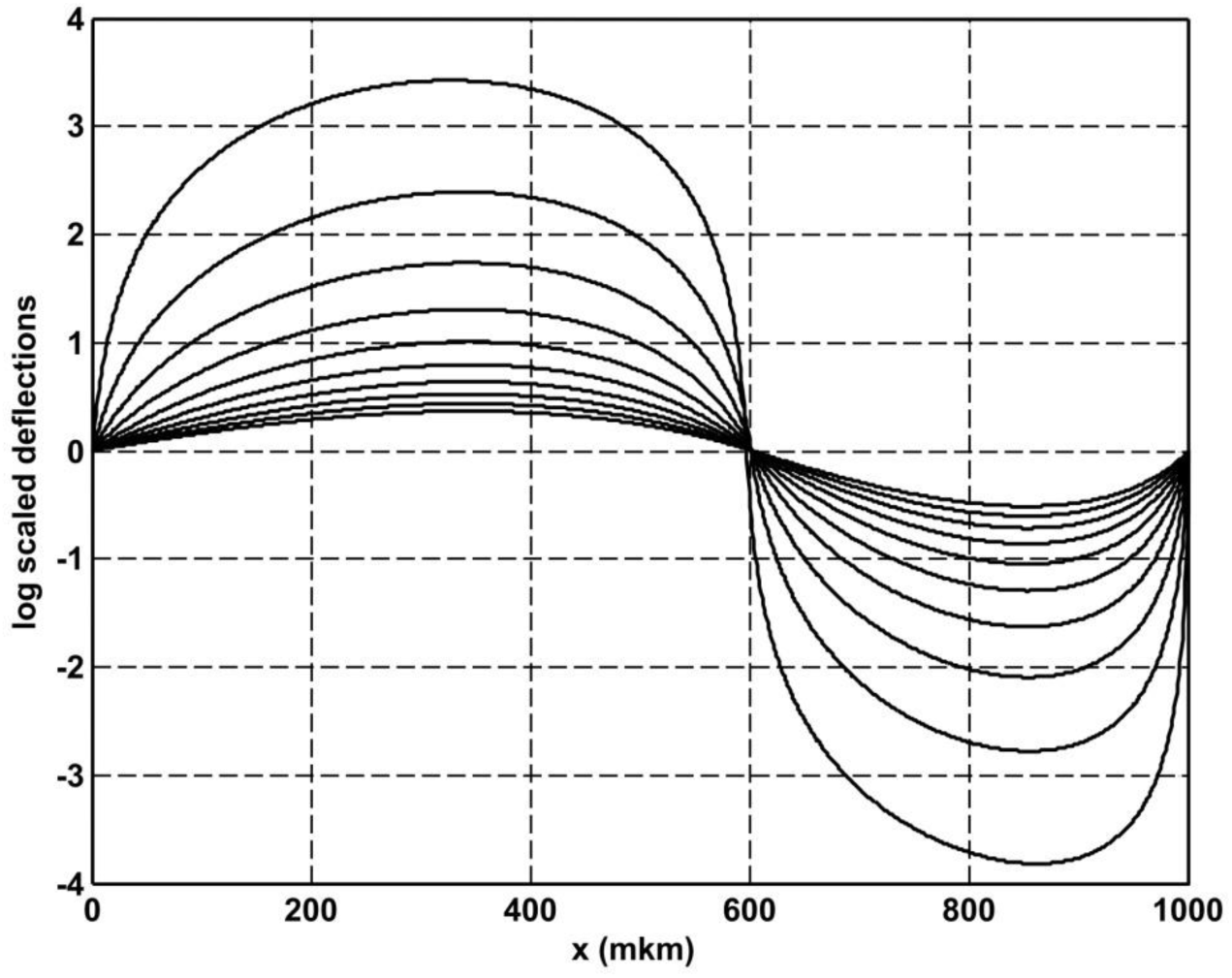
The logarithm scaled membrane deflections for the same set of the linear accelerations Independent of separate or unified scaling of the deflection curves from Figure 4 or Figure 6 we can evaluate the norms 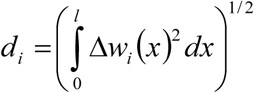 for the raw deflections Δ*w*_*i*_(*x*), *i* =1,…,10. The dependence of the norms *d*_i_ on the acceleration values *γ*_*i*_ gives the estimation of the sensitivity of the utricle membrane as a sensor. This dependence is illustrated by the Figure 7.

**Figure 7.**
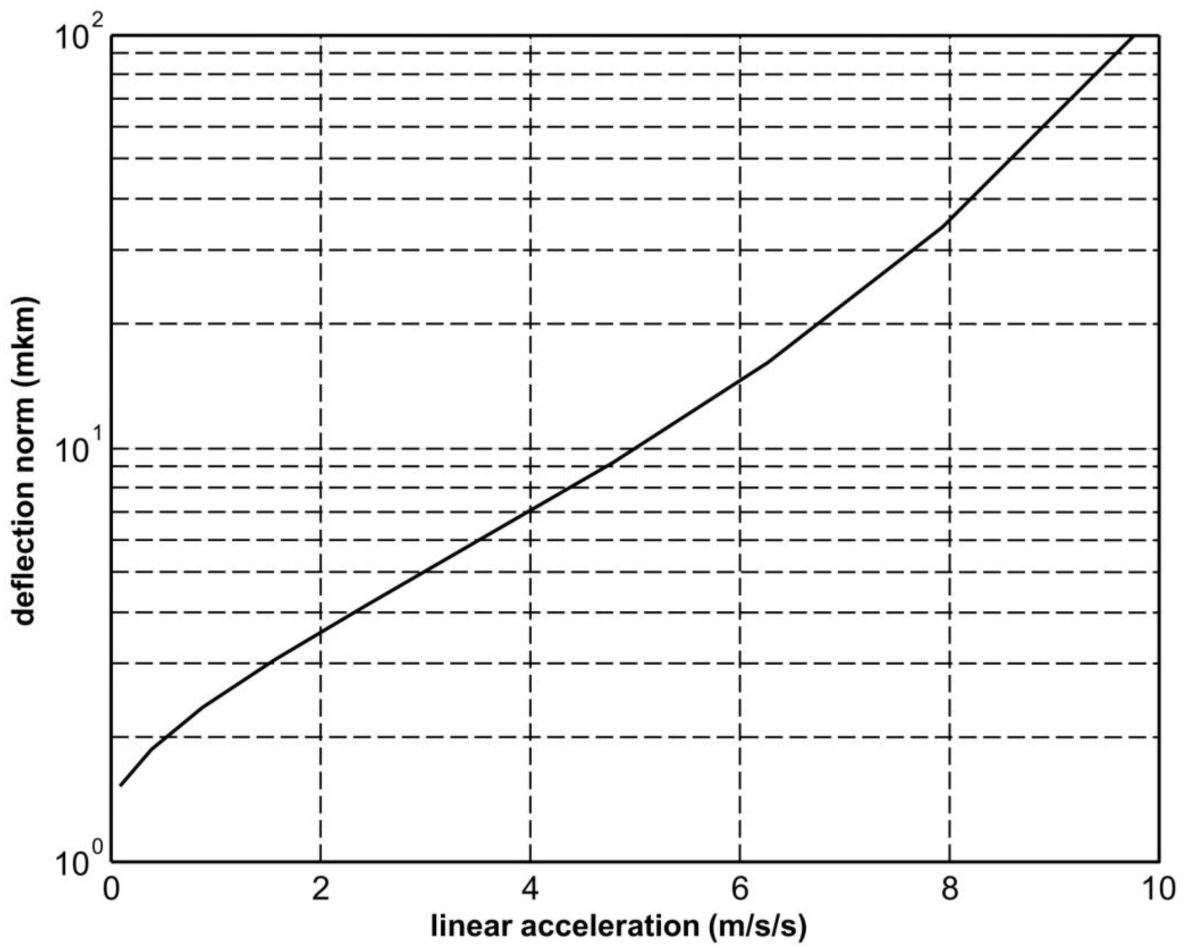
The norms of the membrane deflections for the set of linear accelerations.

## Discussion

### Why are the transversal deflections?

The most basic question immediately arises. Why we need to consider the transversal deflections in spite of the fact that there are already well developed research results supported the longitudinal displacements of otolithic membranes (Hudspeth & Corey, 1977; Hudspeth, Choe, Mehta & Martin, 2000; Colclasure & Holt, 2003). It is also directly demonstrated that the afferent responses of the hair cells can be evoked by longitudinally deflected kinocilia (Eatock, Corey, & Hudspeth, 1987). We are trying to answer this question using numerical results of the proposed model of the utricle membrane deformation under the linear accelerations. The main assumption made in the model is the supposition that the gel like fluid is different from the Kelvin-Voigt fluid (Kluitenberg, 1964) and is simply the conventional viscous fluid with the kinematic viscosity of 1 Poise (Jaeger, 2004.). Only under this supposition the gel layer fluid is able to flow along the membrane from the regions of relatively high elastic pressure to the regions of the lower elastic pressure. The flowing fluid changes the height of the gel layer covering or opening the hair cell bundles, which move in their perforated cavities. From the pictures we can see that only one region divides the utricle membrane in two separate regions: one with increased and another with decreased heights of the gel layer. This region could be associated with the striola. As the deflections are severely dependant on the membrane curvature (Figure 3), the position of the striola is affected by the real membrane curvature, which is not constant in the real 3D membrane shape. The utricle membrane is also submitted to the longitudinal compression and tension. Because of the experimental evidence that the otolith membranes are firmly attached to the bones or more solid tissues (Uzun-Coruhlu, Curthoys & Jones, 2007), they can move longitudinally only by compression or by tension. To estimate the utricle membrane longitudinal displacement we consider again the beam loading scheme in Figure 2. One half of the beam is in the compression whereas another one is in the tension. The maximal beam displacement is observed in the middle section at 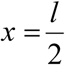. From the regular elasticity problem we can evaluate the maximal displacement *ξ* as:

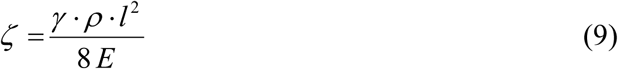

Substituting all available parameters we can find for the linear acceleration *γ* =1 *g* the maximal displacement *ξ* = 0.2817 *mkm*, what is more than10 times less as compared with the transversal deflection using the same parameter set. It is also important to emphasize that this maximal displacement is observed in the middle of the utricle membrane i.e. in the striola region, where the density of the hair cells is reduced. The main question still remains: how can the increased transversal deflections be used to ensure the transduction from mechanical deflections to the electrical activity of the hair cells? The possible answer can be formulated as a hypothesis. It is already found that the gel like liquid is close by its chemical composition to the endolymph and even more is called “mucopolysaccharide gel” (Pandey, 2015). The presence of the polarized mucopolysaccharide molecules lets us hypothesize that the transduction process could be realized on the molecular level similar to transduction in the cochlea inner hair cells (Goussev, 2018) or the semicircular canal hair cells (Gusev, 1994). The principal condition for the molecular transduction is the movement of the polarized molecules in the vicinity of the hair cell bundles. It is exactly this condition that occurs in the utricle membrane with transversal deflections when the hair cell bundles move inside the mesh layer cavities. With the evaluated maximal longitudinal displacements of order of the stereocilia width the hair bundles move mainly up and down in their cavities rather than being bent in the longitudinal direction by the linear acceleration.

### Conclusion

The steady state transversal deflections of the otolithic membranes can be observed along with their longitudinal displacements caused both by the linear acceleration load in the buckling way. As compared with the longitudinal displacements, the transversal deflections are stronger and are concentrated in the regions of maximal density of the hair cells. It is hypothesized that the transduction of transversal deflections to the electrical activity of the hair cells is realized by the molecular mechanisms based on the polarized mucopolysaccharide molecules.

